# Magnetic Resonance Imaging for Improved Brain Tumor Detection

**DOI:** 10.1101/2025.05.06.652446

**Authors:** Blake Benyard, Narayan Datt Soni, Anshuman Swain, Nishi Srivastava, Sunil Kumar Khokhar, Abeer Mathur, Akila Ramesh, Brandon Curl, Arati Desai, Stephen Bagley, Nadir Yehya, Yi Fan, Dushyant Kumar, Ravi Prakash Reddy Nanga, Michael P. Frenneaux, Suyash Mohan, Neil Wilson, Sanjeev Chawla, Ravinder Reddy, Mohammed Haris

## Abstract

Precise demarcation of brain tumor boundaries is critical for optimizing treatment strategies and improving patient outcomes. *In vivo* characterization of tumors using PET/CT and MRI is a clinical standard. PET/CT highlights metabolic processes, while MRI provides information on functional, metabolic, and structural changes. Even with technological advancements in both PET/CT and MRI, a method that can precisely delineate infiltrative tumor boundaries from normal-appearing brain regions (NABR) *in vivo* is still lacking. To address this limitation, we explored a relatively new MR imaging method, the Nuclear Overhauser Effect Magnetization Transfer Ratio (NOE_MTR_), in combination with a gadolinium-based contrast agent (Gd-DOTA) to precisely delineate tumor boundaries in a rat model of infiltrative glioblastoma. NOE_MTR_ imaging was performed in a rat glioma model (n = 5) and in human patients (n = 3) with glioblastoma, both before and after Gd-DOTA administration. The post-Gd-DOTA NOE_MTR_ map was subtracted from the pre-Gd-DOTA map and compared with contrast-enhanced T_1_-weighted images and immunohistological findings. The resulting NOE_MTR_ difference map clearly highlighted both the tumor core and infiltrative boundaries, which were not discernible on the post-contrast T_1_-weighted images. Guided by the NOE_MTR_ difference map, regions of interest (ROI) were drawn to quantify NOE_MTR_ signal changes in the tumor core, tumor boundaries, and NABR post-Gd-DOTA. The tumor core showed a significant ∼43% reduction in NOE_MTR_ signal (p = 0.003), while the tumor periphery exhibited a moderate reduction of ∼10% (p = 0.045). No appreciable change was observed in the NABR (p = 0.371). In contrast, the post-contrast T_1_-weighted signal changes in the tumor core, tumor periphery, and NABR were 33.32% (p = 0.092), 3.8% (p = 0.478), and 8.7% (p = 0.464), respectively. These findings suggest that NOE_MTR_ imaging provides enhanced tumor contrast, particularly at the infiltrative tumor margins, where conventional contrast-enhanced T_1_-weighted MRI may underestimate tumor extent. Histological validation confirmed the presence of infiltrative tumor cells and macrophages in the tumor periphery, as highlighted by the NOE_MTR_ difference map. The NOE_MTR_ difference map showed an extended tumor boundary in the human patient when compared to the T_1_-weighted contrast difference map. Overall, NOE_MTR_ imaging, in combination with Gd-DOTA administration, demonstrates superior delineation of brain tumor boundaries compared to conventional MRI. As NOE_MTR_ imaging is a fast acquisition scan (under 10 minutes) and is performed on a standard 3 Tesla MRI, it can be easily integrated into clinical protocols. By improving the visualization of tumor infiltration and distinguishing tumor regions from NABR, NOE_MTR_ imaging holds promise for advancing neuro-oncological diagnostics and treatment planning.

## Introduction

Accurate tumor identification and precise delineation of brain tumor boundaries are essential for clinicians to tailor interventions, optimize treatment strategies, guide surgical resection while minimizing damage to surrounding healthy tissue, and assess patient prognosis ^1–6^. A multicenter cohort study found that maximal resection of the contrast-enhanced (CE) tumor is associated with improved overall survival in all patients with newly diagnosed glioblastoma (GBM), regardless of age, IDH mutation status, or MGMT (O6-methylguanine-DNA methyltransferase) methylation ^7^. Importantly, in patients younger than 65 years, aggressive resection of both CE and surrounding non-contrast-enhanced (NCE) tumor regions significantly improved overall survival, regardless of tumor subtype ^7^. Another recent study showed that supramaximal resection of NCE tumor regions improved overall survival for all newly diagnosed patients with IDH wild-type GBM^8^. These findings support a revised surgical approach that prioritizes maximal NCE tumor resection in GBM patients, regardless of molecular subtype. This underscores the clear need for precise demarcation of tumor boundaries to enable safe resection while avoiding damage to normal brain regions, which cannot be achieved through random surgical removal of NCE regions.

Although Computed Tomography (CT) and Positron Emission Tomography (PET) using ^18^F-FDG are widely used for tumor diagnosis, they are unable to accurately delineate tumor margins. Moreover, their reliance on ionizing radiation limits their use for repeated monitoring of therapeutic efficacy. In contrast, conventional Magnetic Resonance Imaging (MRI) does not involve radiation and can be safely repeated without adverse effects, making it more suitable for ongoing assessment and treatment planning. However, conventional magnetic resonance imaging (MRI) techniques often face challenges in distinguishing tumor margins due to the complex and heterogeneous nature of brain tumors ^9–12^. In particular, the transition between tumor tissue and normal-appearing brain regions (NABR) can be subtle, lacking well-defined edges, which complicates treatment planning ^13^.

To date, contrast-enhanced T_1_-weighted MRI remains the gold standard for visualizing brain tumors and differentiating them from NABR ^14^. This technique relies on the accumulation of contrast agents, such as gadolinium-based compounds, in the extracellular-extravascular space (EES) of tumor tissue, thereby enhancing visualization of abnormal regions. Despite its widespread use, several limitations persist, including insufficient contrast between the tumor and peritumoral regions, partial volume effects that obscure finer structural details, and difficulty detecting tumor infiltration into the surrounding brain tissue ^14^. These challenges are particularly pronounced in infiltrative tumors, where malignant cells may extend beyond the visibly enhanced regions, leading to potential underestimation of tumor extent and suboptimal treatment planning.

To address these limitations, Nuclear Overhauser Effect (NOE)-weighted MRI has emerged as a promising alternative, offering enhanced sensitivity to molecular and structural changes in the tumor microenvironment ^15–28^. This advanced imaging technique uses indirect detection of membrane lipids and other macromolecular interactions to provide contrast based on biophysical properties. Given that tumor cells often exhibit compromised plasma membrane integrity, NOE-mediated magnetization transfer ratio (NOE_MTR_) imaging has the potential to improve discrimination between tumor tissue and NABR ^23^.

In this study, we explored the potential of NOE_MTR_ imaging in conjunction with gadoterate meglumine (Gd-DOTA) administration to enhance tumor contrast and improve visualization of tumor boundaries. By integrating NOE_MTR_ with conventional contrast-enhanced MRI, we aim to capture a more comprehensive representation of tumor extent, potentially overcoming the limitations of standard imaging methods in delineating tumor boundaries.

## Materials and Methods

We have performed NOE_MTR_ imaging and conventional imaging on both a rodent model of glioblastoma and human patients with recurrent glioblastoma. The animal experimental protocols used in this study were approved by the Institutional Animal Care and Use Committees of the University of Pennsylvania. The study was conducted in accordance with an approved Institutional Review Board protocol from the University of Pennsylvania.

### Rodent brain tumor model

Five Sprague-Dawley rats (∼6 weeks old, ∼140 g) were used to generate a glioma, tumor-bearing model. Briefly, rats were anesthetized with isoflurane, and their scalp hair was removed using a hair trimmer and hair removal cream. Rats were positioned with their heads fixed on the stereotaxic apparatus equipped for a continuous supply of isoflurane to induce a surgical plane of anesthesia. Bupivacaine (under the scalp; 2 mg/kg) and meloxicam (subcutaneous; 2 mg/kg) were administered as analgesics. After sterilizing the scalp, an incision was made to expose the skull. Following stereotaxic navigation to 3 mm lateral (right) and 3 mm posterior to bregma, a burr hole was drilled to inject a 5 μl suspension containing ∼50,000 glioblastoma (F98) cells in phosphate buffer into the cerebral cortex (2 mm deep) using a Hamilton syringe and a 32-gauge needle. These rats underwent MR imaging three weeks after tumor cell implantation.

### 9.4T MR Imaging

MRI was performed on isoflurane (1.5%) anesthetized rats using a 9.4T horizontal bore preclinical MR scanner (Bruker) and a 30-mm diameter proton volume coil (m2m Imaging Corp., OH). Animals were kept under anesthesia (1.5% isoflurane in 1 liter/min oxygen), and their body temperature was maintained by circulating air generated and blown through a heater (SA Instruments, Inc., Stony Brook, NY). Respiration and body temperature were continuously monitored using an MRI-compatible small animal monitoring system (SA Instruments, Inc., Stony Brook, NY).

A localizer was acquired first, followed by a T_1_-weighted FLASH (TE= 4 ms , TR = 498 ms, four averages, 16 slices) and T2-weighted RARE (TE1/TE2 = 33/121 ms, TR = 3s, two averages, 16 slices, rare factor = 6. For NOE MRI acquisitions, a water saturation shift referencing (WASSR) image (TE = 4 ms, TR = 410 ms, 22 frequency offsets from 0 to 1 ppm in steps of 0.1 ppm, B_1rms_ = 0.1 μT) was acquired for correcting B_0_ inhomogeneities ^29^. A full Z-spectrum was acquired for the saturated images with 52 offsets from +5 to -5 ppm in steps of 0.2 ppm). The acquisition parameters were: TE = 4, TR = 6s, B_1_ = 1.0 μT, saturation duration (tsat) = 3 s, and two averages. An unsaturated image (with the same parameters as the saturated images), was additionally acquired. The FOV was 20 mm x 20 mm with an image matrix size of 128 × 128, resulting in an in-plane resolution of 0.156 mm x 0.156 mm for all images.

An intravenous injection (IV) of a gadolinium-based contrast agent (Gadoterate meglumine, Dotarem, France; 0.1 mmol/kg) (Gd-DOTA) was administered through a pre-inserted catheter in the tail vein. To monitor changes in signal intensity, T_1_-weighted images and Z-spectrum data were acquired again on the same slice 30 minutes post-injection. After imaging, the rats were perfused, and the brains were extracted for hematoxylin staining. The total scanning time was ∼90 minutes (**Fig. 1**).

**Figure 1:**
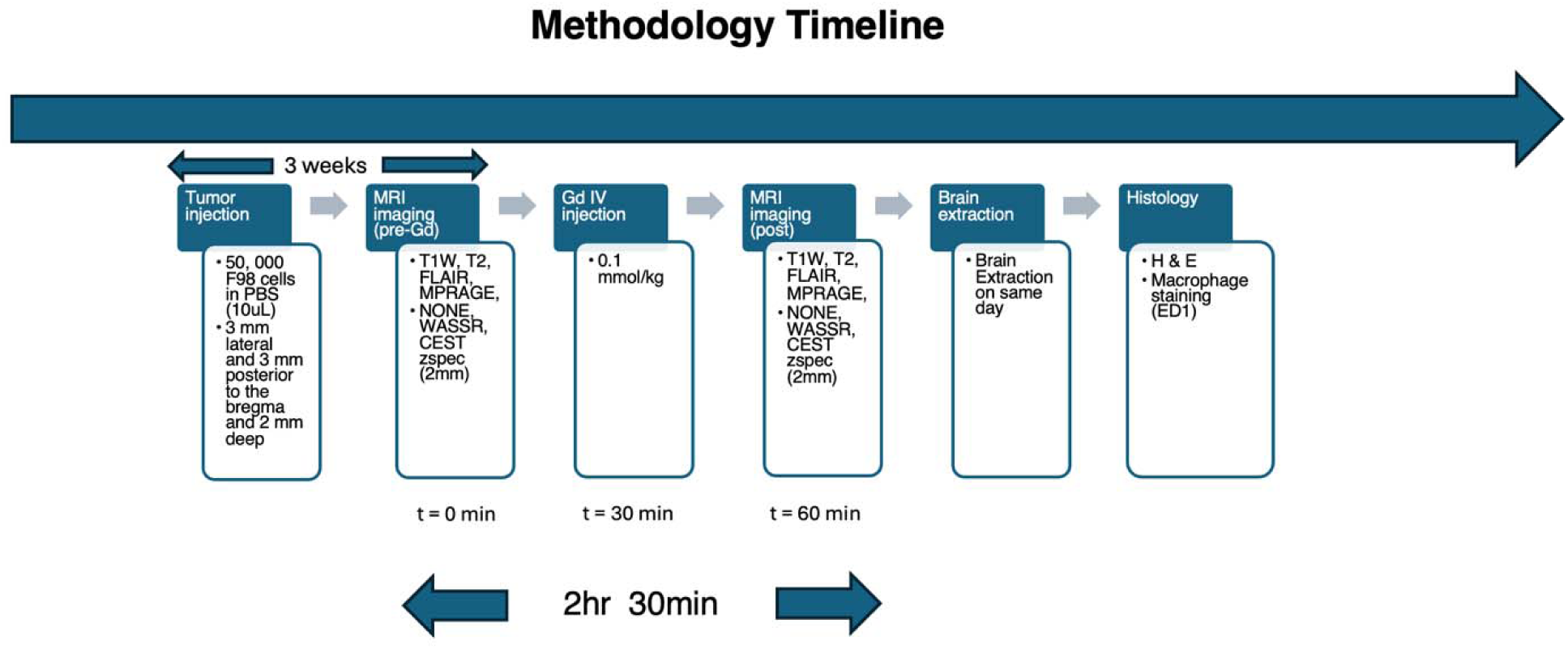
A timeline description of the methodology of this study. Rats were injected with the tumor cells (F98) and scanned three weeks later. Pre- and post-Gd imaging was performed using the same MR sequences (T_1_, T_2_, CEST, NOE). The brains of these rats were extracted the same day for histological analysis (hematoxylin and ED1 staining).

### Immunohistochemistry

After dissection and fixation for 24 hrs in 4% cold paraformaldehyde, rat brains were washed in 1X PBS and processed through a series of alcohols, xylenes, and wax for 14 hours, followed by embedding in paraffin. Thin 5 µm paraffin coronal sections of the brain were collected from the tumor site and mounted onto the slides. After drying overnight, the slides were baked at 60°C for 20 min, then rehydrated in xylenes, graded alcohols, and water. For antigen retrieval, slides were then exposed to a 60°C wash with 1X citrate buffer (#H-3300; Vector Labs) for 30 min, and, following rinses in PBS, they were incubated at 37°C in 20 µg/ml Proteinase K in 1xPBS and treated with a 15 min wash in 3% H_2_O_2_ to block endogenous peroxidase.

Slides were placed in a humid chamber with 3% bovine serum albumin (1X PBS) for 1 hr at room temperature. Each section was treated with a 1:500 dilution of Mouse-anti-rat ED1 (CD68) (BioRad # MCA341GA) overnight at 4°C in a humid chamber. After washing in 1X PBS, sections were treated with horse-anti-mouse biotinylated antibodies (Vector Labs, Newark, CA) for 30 min using the peroxidase ABC kit (PK-6200) according to the manufacturer’s instructions. After washing in 1X PBS, the sections were treated with chromogenic immPACT DAB (Vector Labs; SK-4100). This was followed by washing in ddH_2_O and counterstaining with hematoxylin. The sections were dehydrated in a graded series of alcohols and xylene and then mounted with Permount.

### Human patient imaging

Three subjects with recurrent glioblastoma were recruited for scanning. All imaging was performed on a 3T whole-body scanner (MAGNETOM Prisma, Siemens Healthcare, Erlangen, Germany) with a body transmit/32-channel receive proton phased-array head coil. An intravenous injection (IV) of a gadolinium-based contrast agent (Gadoterate meglumine, Dotarem, France; 0.1 mmol/kg) (Gd-DOTA) was administered. T_1_-weighted images and canonical Z-spectrum data were acquired pre- and post-administration of Gd-DOTA. The CEST acquisition parameters include number-of-slices = 1, slice thickness = 10 mm, in-plane resolution = 1 x 1 x 10 mm^3^, matrix size = 240 x 196, gradient echo readout TR = 3.5 ms, TE = 1.79 ms, short TR = 6s, read-out flip-angle = 10°, averages = 2, and saturation pulse of B_1,rms_ = 0.62 µT with a saturation length of 3 s. The CEST-weighted images were acquired at varying saturation offset frequencies +5 to -5 ppm in steps of 0.25 ppm, and ± 100 ppm) relative to the water resonance, with a step size of 0.25 ppm. The total scanning time, including NOE imaging, was under 60 minutes.

To further demonstrate that NOE_MTR_ can delineate the infiltrative tumor boundary, the subtracted NOE_MTR_ map (pre-vs post-Gd) from one of the patients was co-registered, overlaid on the post-Gd T_1_-weighted image, and integrated into Brainlab (**supplementary figure 1**), a neuro-navigation system routinely used in clinic for tumor surgical guidance ^30^.

### Data Processing

All image datasets will be transferred to a dedicated workstation for image processing. Image processing, including the generation of NOE_MTR_ maps, pixel-by-pixel co-registration of parametric maps with anatomical images, and quantitative analyses, will be performed using in-house-developed algorithms written in MATLAB (version R2026b), as described previously^31^. Acquired images will be corrected for B_0_ and B_1_ inhomogeneities, and NOE_MTR_ contrast will be calculated from equations [1]. NOE_MTR_ contrast will be mapped onto the anatomical image as false colors.

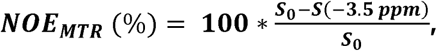

where S_0_ is the signal intensity without saturation or with a saturation pulse applied at 100 ppm (NONE). To delineate changes in tissue contrast, the post-Gd-DOTA NOE_MTR_ map was subtracted, pixel-by-pixel, from the pre-Gd-DOTA NOE_MTR_ map.

## Results

Figure 2 (also **supplementary figure 2**) shows NOE_MTR_ maps from a rat GBM model before (**Fig. 2a**) and after (**Fig. 2b**) Gd-DOTA administration, along with the calculated difference map (**Fig. 2c**), highlighting areas with the most pronounced contrast changes. The corresponding T_1_-weighted images (pre-contrast and post-Gd-DOTA) and the calculated difference maps are shown in **Fig. 2d–f** for comparison. The NOE_MTR_ difference map (**Fig. 2c**) reveals a clear reduction in NOE_MTR_ signal within both the tumor core and tumor periphery, while regions corresponding to the NABR exhibit no appreciable changes. Notably, no visible signal change was observed in the tumor periphery on the post-contrast T_1_-weighted image. However, overlaying the NOE_MTR_ difference map onto the post-contrast T_1_-weighted image clearly demonstrates tumor boundaries beyond those enhanced on conventional post-contrast T_1_-weighted imaging. This superior delineation underscores the ability of NOE_MTR_ imaging to capture molecular-scale variations that may be missed by traditional contrast-enhanced MRI.

**Figure 2:**
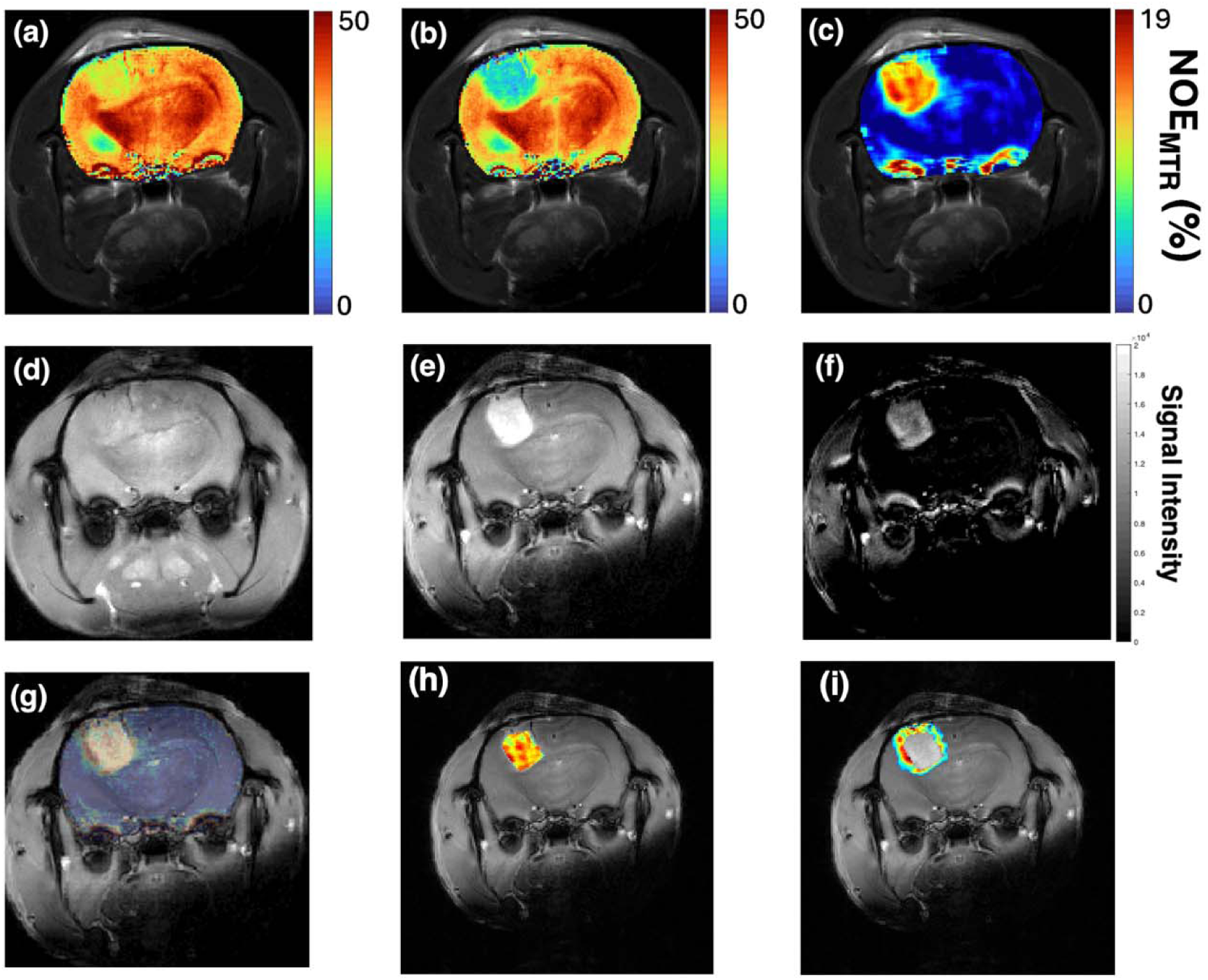
*In vivo* NOE_MTR_ maps from a brain tumor-bearing rat brain before and after the infusion of gadoterate meglumine (Gd-DOTA). Image (a) displays the pre-infusion NOE_MTR_ map, while image (b) shows the post-infusion NOE_MTR_ map. Image (c) illustrates the subtraction of the post-infusion NOE_MTR_ map from the pre-infusion NOE_MTR_ map, and image (d-f) displays the corresponding T_1_-weighted pre-Gd, post-Gd, and difference map. The overlayed difference NOE_MTR_ map on the post-Gd T_1_-weighted image (g), NOE difference map in the tumor core (h), and difference map outside the tumor core (i) highlights distinct tumor boundaries as compared to the post-Gd T_1_-weighted image.

To further validate these findings, histological analysis was performed. **Figure 3** presents hematoxylin-eosin and ED1-stained tumor sections, revealing the spatial distribution of the tumor core and tumor boundary. Zoomed-in regions from the tumor boundary, where post-Gd-DOTA NOE_MTR_ signal changes were observed, correspond to macrophage-rich regions containing infiltrative tumor cells, suggesting a biologically relevant transition zone between tumor tissue and adjacent normal brain (**Fig. 4**). Notably, the zoomed-in tumor region shows no signal enhancement on the post-contrast T_1_-weighted image at the tumor boundary (**Fig. 4**). The bar graphs illustrate a quantitative comparison of NOE_MTR_ contrast across distinct tissue compartments. The bar plot summarizes average NOE_MTR_ values in the tumor core, tumor periphery, and contralateral NABR across all rats (n = 5) (**Fig. 4**). The tumor core demonstrates the most pronounced NOE_MTR_ signal drop following Gd-DOTA administration, while the peritumoral region shows a moderate yet distinguishable decrease. In contrast, NOE_MTR_ values in the NABR remain largely unchanged, underscoring the specificity of the observed alterations in contrast within tumor-associated regions.

**Figure 3:**
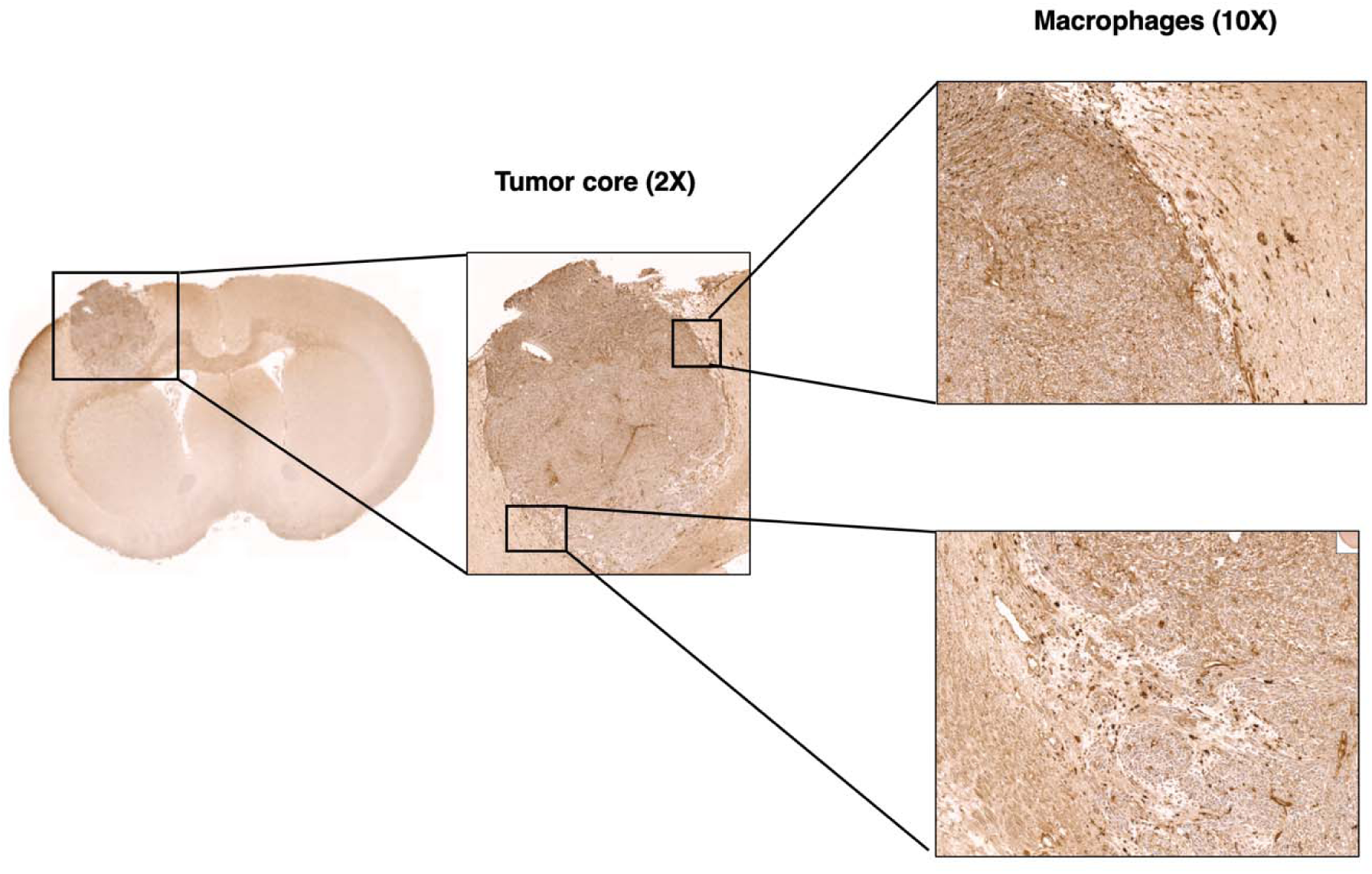
Immunohistochemical staining was performed using Hematoxylin and Anti-CD68 antibody [ED1] to visualize general tissue architecture and macrophage distribution, respectively. The hematoxylin-stained section shows the full brain slice corresponding to the tumor region from one representative rat, with zoomed-in views highlighting the tumor core, tumor boundary, and regions enriched with macrophages. ED1 staining confirms the presence of macrophages, particularly concentrated along the tumor periphery.

**Figure 4:**
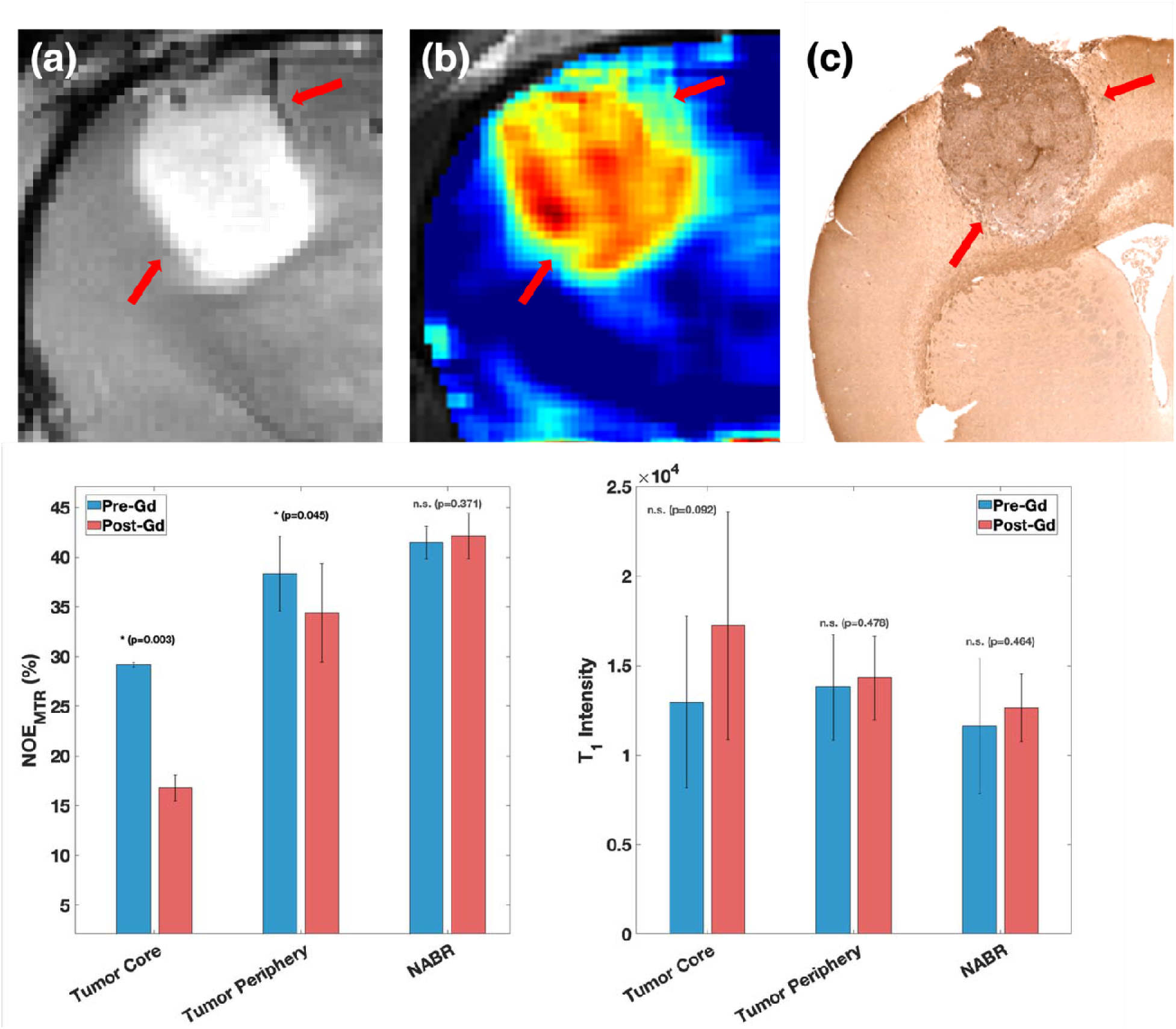
The zoomed-in tumor area shows clear uptake of Gd-DOTA contrast in the tumor on the post-contrast T_1_-weighted image (a), but it does not demarcate the tumor boundary (red arrows). In contrast, the NOR_MTR_ difference map (b) highlights tumor regions extending beyond the signal observed on the post-contrast T_1_-weighted image, which was further confirmed by the immunohistochemistry. The immunohistochemistry image clearly shows infiltrated tumor cells and macrophages at the tumor periphery (red arrows). Bar plot representing average NOE_MTR_ percentages across regions of interest (ROIs) in the tumor core, tumor periphery, and contralateral normal-appearing brain region (NABR), measured pre- and post-Gd-DOTA infusion (n=5). A decrease in NOE contrast is observed in both the tumor core and tumor periphery, while no appreciable change is observed in NABR.

Quantitative analysis supports these visual observations. Following Gd-DOTA administration, the tumor core exhibited the largest percentage decrease in NOE_MTR_ signal (−42.5%, p = 0.003), with an average drop from 29.2 ± 0.2 to 16.8 ± 1.3. The tumor periphery demonstrated a moderate reduction (−10.3%, p = 0.045) in NOE_MTR_ signal (from 38.3 ± 3.7 to 34.4 ± 5.0). In contrast, the NABR showed a negligible change in NOE_MTR_ signal (+1.6%, p = 0.371), with values remaining stable (from 41.4 ± 1.7 to 42.1 ± 2.3). These findings emphasize the specificity of the NOE_MTR_ signal drop within tumor-associated tissue compartments.

In comparison, changes observed on T_1_-weighted images were more subtle and less spatially specific. The tumor core exhibited a 33.2% increase in T_1_-weighted signal (from 12,956 ± 4,797 to 17,251 ± 6,361, p = 0.092), while the tumor periphery showed a 3.8% increase (from 13,788 ± 2,922 to 14,310 ± 2,336, p = 0.478) and the NABR regions showed an 8.7% increase in T_1_-weighted signal (from 11,618 ± 3,764 to 12,632 ± 1,905, p = 0.464), indicating statistically insignificant contrast differentiation between normal and tumor-adjacent tissues. These results highlight the enhanced sensitivity and spatial specificity of NOE_MTR_ over conventional T_1_-weighted imaging in detecting subtle molecular alterations in and around the tumor.

**Figure 5** shows the T_1_-weighted and NOE_MTR_ maps pre- and post-Gd contrast from a human patient with glioblastoma. The NOE_MTR_ difference map clearly highlights the extended tumor boundary (arrows) compared with either the post-contrast T_1_-weighted image or the T_1_-weighted difference image. Another patient with recurrent GBM shows an extended tumor boundary on the subtracted NOE_MTR_ map compared with the post-contrast T_1_-weighted image (**Fig. 6**). The subtracted NOE_MTR_ map, overlaid on the post-Gd T_1_w images, reveals a larger area of tumor infiltration (**Fig. 7**). At one month of follow-up, the patient exhibited tumor growth in the same regions as depicted on the subtracted NOE_MTR_ map at baseline (**Fig. 7**). The subtracted NOE_MTR_ map (pre-vs post-Gd) was co-registered and overlaid on the post-Gd T_1_-weighted image and integrated into the neuro-navigation system for tumor tissue excision (**Fig. 7**). The subtracted NOE_MTR_ map guided excision of tumor tissue extending beyond the contrast-enhancing border showed infiltrative tumor cells on histopathological analysis and immunohistochemistry (IHC) for Ki67.

**Figure 5:**
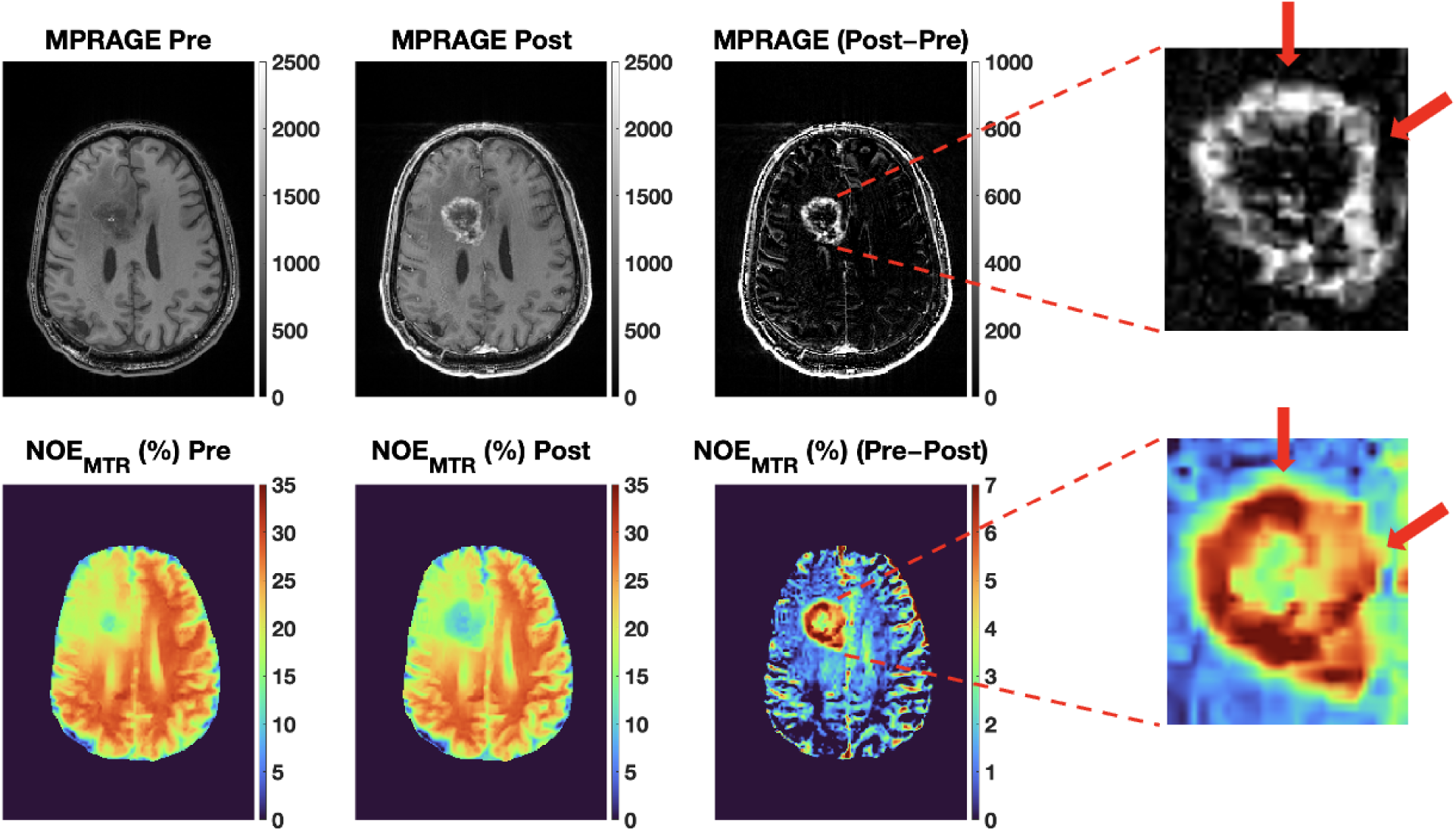
The first row is MPRAGE (T1-weighted) images (pre-Gd, post-Gd, and the difference map) of a human participant with a glioblastoma at 3T. The second row is the respective NOE_MTR_ maps. The zoomed-in figure shows how the NOE_MTR_ map highlights the tumor boundary on the T1-weighted difference image and the NOE_MTR_ difference map. Red arrows are showing the extended tumor boundary on NOE_MTR_ difference map.

**Figure 6:**
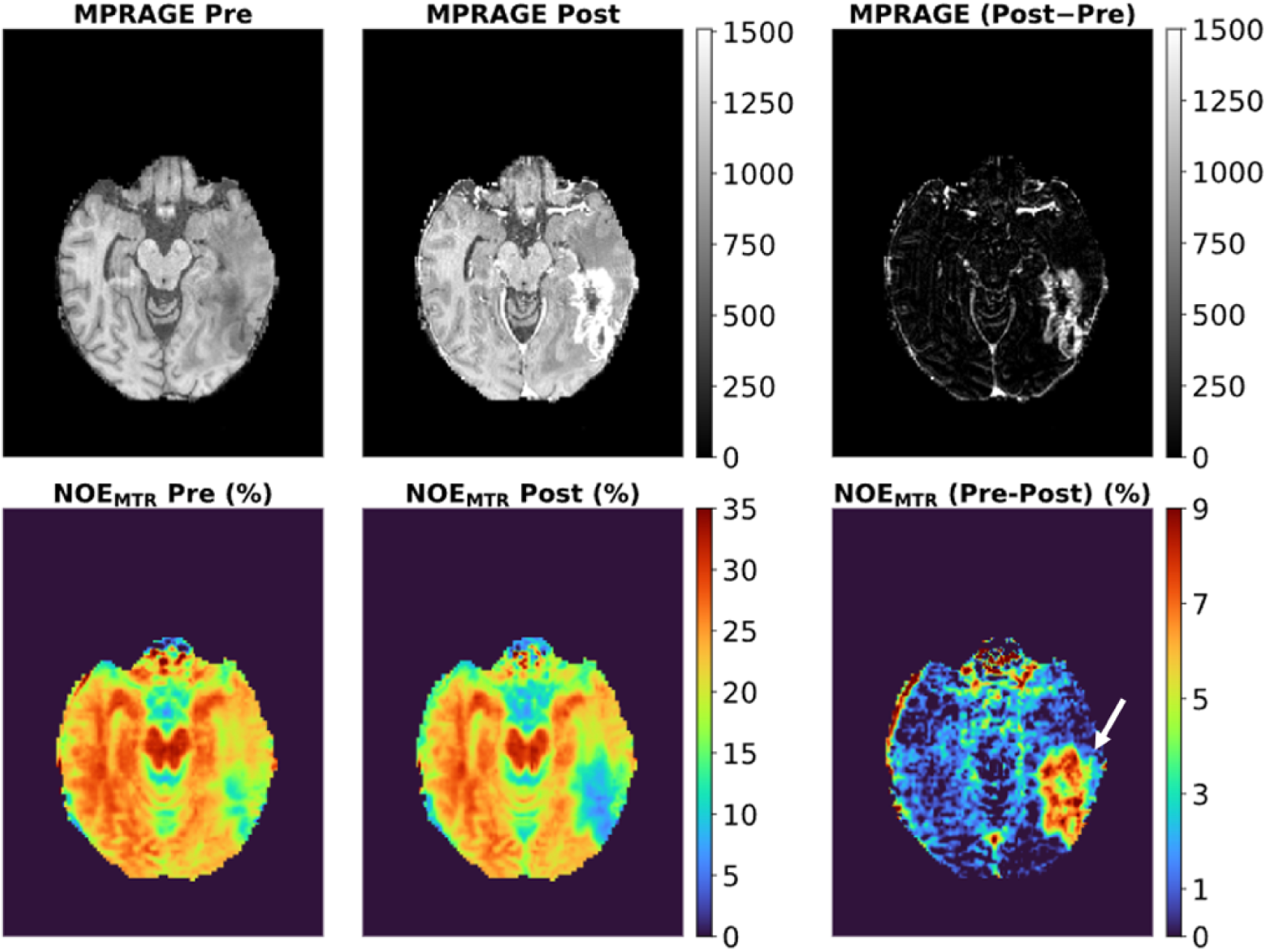
The first row shows T1-weighted images (pre- and post-Gd, and the subtracted map) from a GBM patient scanned on 3T MRI. The second row shows the respective NOEMTR maps. The subtracted NOEMTR map shows the extended tumor boundary beyond the contrast enhancement (white arrow).

**Figure 7:**
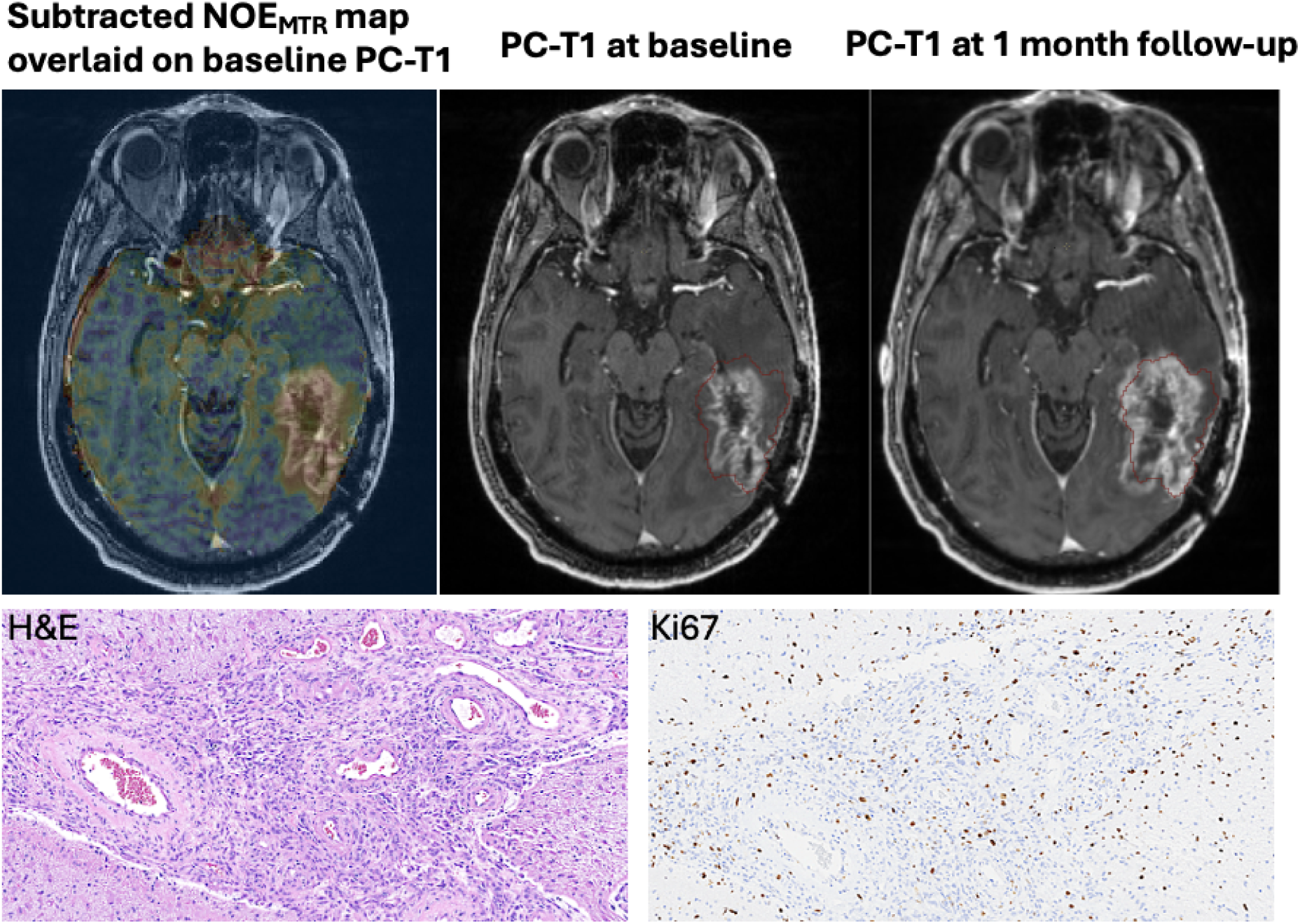
(same patient as shown in Figure 7). The subtracted NOE_MTR_ map (pre-vs post-Gd) overlaid on the baseline post-Gd T1w image shows a larger area of tumor infiltration. The post-Gd T1w image at 1-month follow-up shows tumor progression in regions similar to those highlighted on the subtracted NOE_MTR_ map at baseline. This was further confirmed by histopathology (H&E) and immunohistochemistry (IHC; Ki-67). These findings suggest that tumor abnormalities visible on the subtracted NOE_MTR_ map extending beyond the contrast-enhancing region can predict future tumor progression.

## Discussion

Our results demonstrate a marked improvement in tumor boundary visualization using NOE_MTR_ imaging following Gd-DOTA administration. ROI analysis revealed significant contrast differences across tumor regions, with a mean signal reduction of (∼13%) in the tumor core and (∼4%) in the tumor periphery, while the NABR remained largely unchanged (∼1%). These findings underscore the sensitivity of NOE_MTR_ imaging to molecular interactions and alterations in macromolecular content within tumor regions.

NOE_MTR_ is more sensitive in depicting the tumor boundary than post-contrast T_1_-weighted imaging. As shown in the bar graph, although the change in T_1_-weighted signal intensity is not significant, it results in a significant change in NOE_MTR_ contrast in both the tumor core and tumor boundary. While there is a slight, non-significant increase in post-contrast T_1_-weighted signal intensity within the NABR, it does not result in a noticeable change in NOE_MTR_ contrast. This is likely because, in NABR, the T_1_-weighted signal change is primarily due to the shortening of water proton T_1_ in the circulating blood, which does not have appreciable cross-relaxation effects with the protons on lipids or macromolecules in NABR.

In contrast, both the tumor core and boundary exhibit BBB leakage, allowing Gd-DOTA molecules to accumulate. This accumulation leads to T_1_ shortening of water protons that are in direct contact with tumor cell membrane lipids and macromolecules. The significant cross-relaxation between these water protons and the aliphatic protons of lipids and macromolecules exposed in cancerous cells contributes to the observed decrease in NOE_MTR_ contrast. The loss of ECM structure in the tumor core/periphery results in more mobile lipids and macromolecules, which reduce cross-relaxation between water protons and aliphatic protons on these molecules, thereby reducing the NOE_MTR_ effect. Since the disintegration of extracellular matrix structure is significantly higher in the tumor core than in the periphery, the reduction in NOE_MTR_ is higher than that in the periphery. Notably, the NOE_MTR_ difference maps aligned with histological regions of tumor infiltration, revealing a biological sensitivity that may be obscured in conventional MR imaging. Histological validation reveals strong spatial correspondence between NOE_MTR_ difference maps and the distributions of tumor cells and macrophages. This suggests that NOE_MTR_ imaging adds potential value for characterizing tumor heterogeneity and guiding targeted therapies. Furthermore, NOE_MTR_’s minimal variation in NABR signal post-Gd contrast highlights its potential for reliable tumor-to-background discrimination in both pre- and post-Gd settings.

PET/CT is one of the most widely used imaging modalities for characterizing tumors in clinical practice. A recent study highlighted that the widespread use of CT scans has raised significant public health concerns due to the potential for radiation-induced cancers. Although children are more sensitive to radiation and face a higher risk per scan, adults receive most CT exams, contributing to most of the projected future cancer cases, particularly lung, colon, leukemia, and bladder cancers ^32^. If current usage and radiation dose practices continue, CT-related cancers could eventually comprise up to 5% of all new cancer diagnoses annually ^32^. This risk may be further exacerbated by the increasing use of PET/CT scans, which combine two imaging modalities and deliver significantly higher radiation doses than standard CT alone ^32–34^, thereby , amplifying cumulative exposure and the associated cancer risk. These findings underscore the urgent need to reduce unnecessary imaging and to optimize radiation doses across all modalities. In this context, MRI with non-ionizing radiation and improved capabilities for precise brain tumor delineation could serve as a safer alternative for certain clinical applications.

Since clinical imaging protocols for brain tumors already incorporate contrast-enhanced imaging, integrating NOE_MTR_ into the existing workflow presents a valuable opportunity to enhance tumor characterization without necessitating significant changes to current clinical practice. As a proof of concept, we performed NOE_MTR_ imaging in a patient with glioblastoma, which clearly highlighted the extended tumor boundary not depicted on the conventional post-contrast T1-weighted image and was further validated by histopathology and immunohistochemistry. The demarcation of the tumor lesion by the NOE_MTR_-subtracted map is not only seen in the contrast-enhancing tumor but also in a patient with recurrent glioblastoma who shows no enhancement on post-Gd contrast, while the NOE_MTR_ difference map clearly demarcates the tumor lesion (**Supplementary Figure 3).** By leveraging the additional contrast provided by NOE_MTR,_ particularly in regions of tumor infiltration that may appear normal on conventional imaging, this approach can improve the sensitivity and specificity of tumor boundary delineation. The ability to detect subtle changes in tissue microstructure and to differentiate tumor infiltration from that of surrounding normal-appearing brain regions makes NOE_MTR_ a promising adjunct to standard contrast-enhanced MRI. Ultimately, incorporating NOE_MTR_ into routine clinical imaging could enable more accurate diagnosis, facilitate more precise treatment planning, and better monitor therapeutic response over time. While we have demonstrated the utility of this approach in characterizing a glioblastoma tumor, its underlying principles and advantages are broadly applicable to the clinical imaging of a wide range of solid tumors.

Given its improved differentiation of tumor regions, NOE_MTR_ imaging emerges as a promising alternative to existing conventional imaging methods for comprehensive tumor characterization. For proof-of-principle purposes, we employed a single slice using a two-dimensional imaging sequence. However, implementing a three-dimensional acquisition of NOE_MTR_ acquisition to enable complete brain tumor mapping would be relatively straightforward.

## Conclusion

Our findings suggest that NOE_MTR_ imaging, particularly in combination with Gd-DOTA administration, offers a significant advantage over conventional T_1_-weighted MRI in delineating tumor boundaries with greater precision. By providing enhanced molecular contrast, NOE_MTR_ imaging enables improved differentiation among the tumor core, peritumoral infiltration, and surrounding normal brain tissue. This innovative approach has the potential to refine treatment planning, optimize surgical resection, and improve patient outcomes in oncology.

## Supplementary Information

Supplementary Figure 1 shows the subtracted NOE_MTR_ map, registered to the post-contrast T_1_-w image, and integrated into a neuro-navigation system. The subtracted NOE_MTR_ map guided the surgical excision of tumor tissue extending beyond the contrast-enhancing border, a finding further validated by histopathology. This finding suggests that NOE_MTR_ imaging in combination with Gd-DOTA administration offers a significant advantage over conventional MRI in delineating tumor boundaries. Supplementary Figure 2 demonstrates the clear differences in NOE_MTR_ contrast pre- and post-Gd-DOTA administration from a rat brain GBM model. The corresponding NOE_MTR_ difference map provides better delineation of the tumor boundary than the post-Gd T_1_-weighted image, further validating the technique’s ability to better demarcate the true tumor infiltrative boundary. Supplementary Figure 3 shows a patient with recurrent glioblastoma with no enhancement on post-Gd T_1_-weighted images or on the difference T_1_-weighted image. However, the difference in the NOE_MTR_ map clearly highlights the tumor regions. This pronounced signal reduction suggests that NOE_MTR_ imaging is more sensitive to molecular alterations induced by the Gd-DOTA contrast agent, likely due to cross-relaxation between water and macromolecules.

**Supplementary Figure 1:**
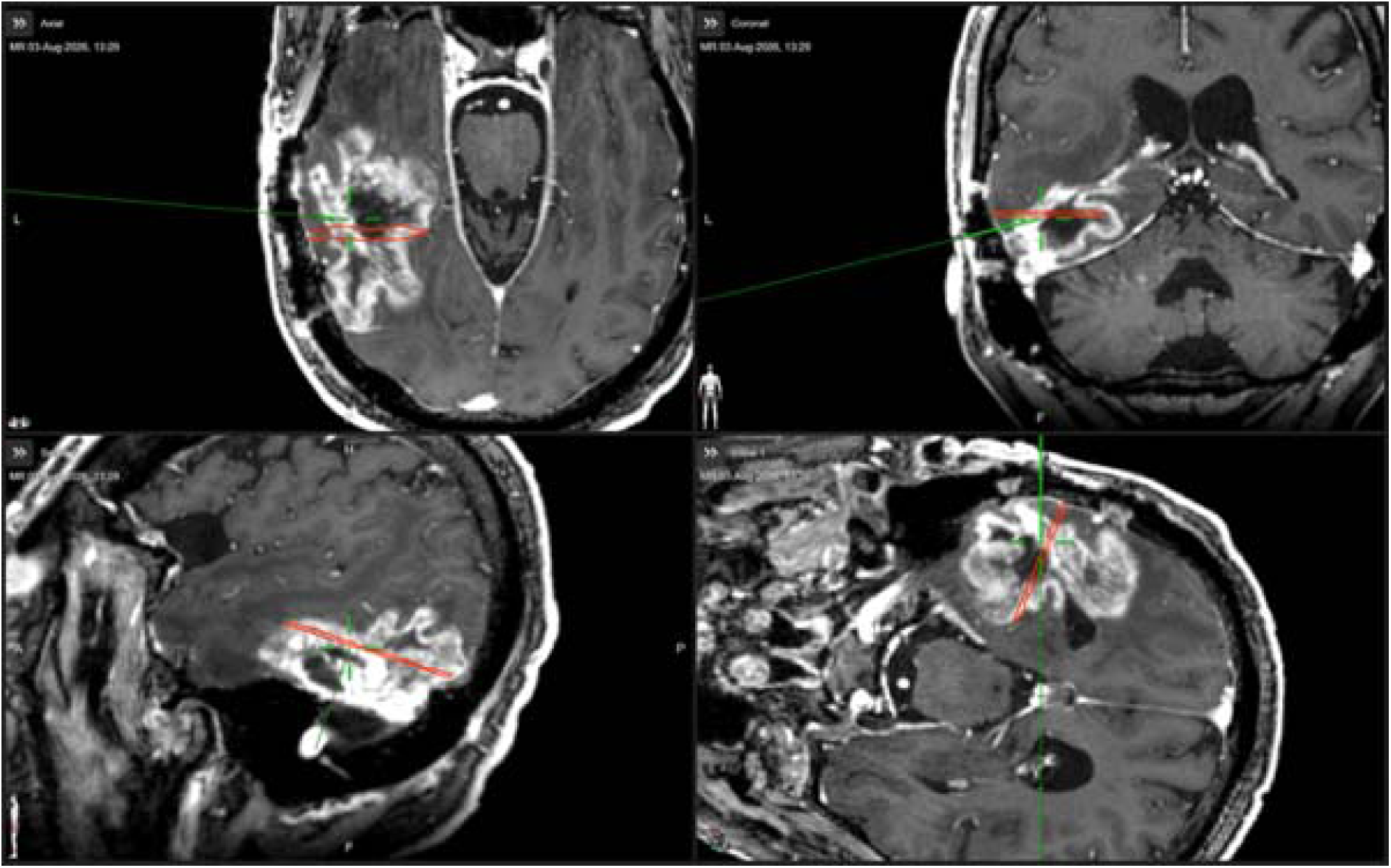
The subtracted NOE_MTR_ map (pre-vs post-Gd) co-registered and overlaid on the post-Gd T_1_-weighted image and integrated into the neuro-navigation system for tumor tissue excision, as guided by tumor abnormality visible on the subtracted NOE_MTR_ map for histopathological confirmation.

**Supplementary Figure 2:**
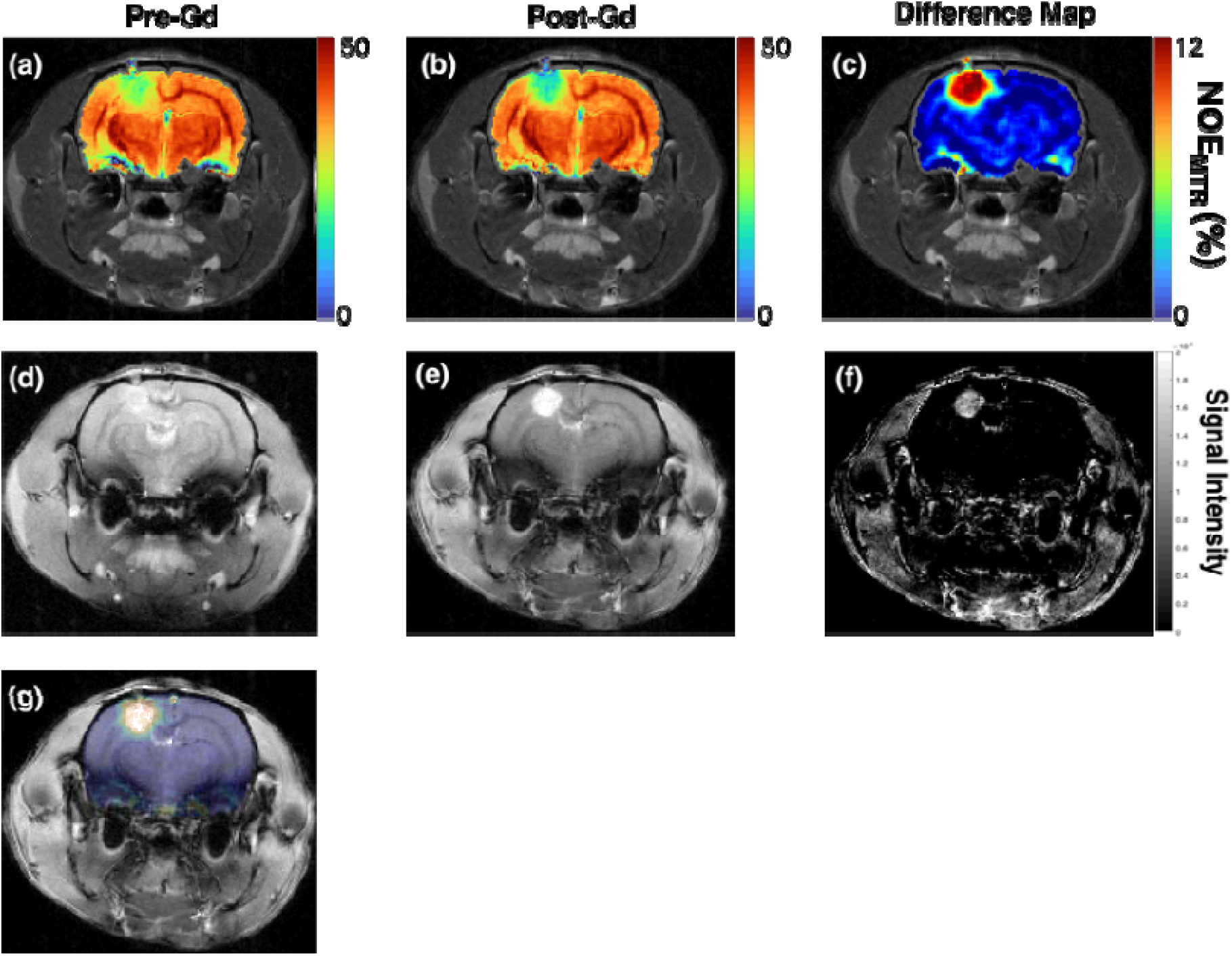
In vivo NOE_MTR_ maps from the brain tumor-bearing rat brain in Figure 5 before and after the infusion of gadoterate meglumine (Gd-DOTA). Image (a) displays the pre-infusion NOE_MTR_ map, while image (b) shows the post-infusion NOE_MTR_ map. Image (c) illustrates the subtraction of the pre-infusion NOE_MTR_ map from the post-infusion NOE_MTR_ map, and image (d-f) displays the corresponding T_1_-weighted pre-Gd, post-Gd, and difference map. The subtracted NOE_MTR_ map (g) highlights distinct tumor boundaries compared with the post-Gd T_1_-weighted image.

**Supplementary figure 3:**
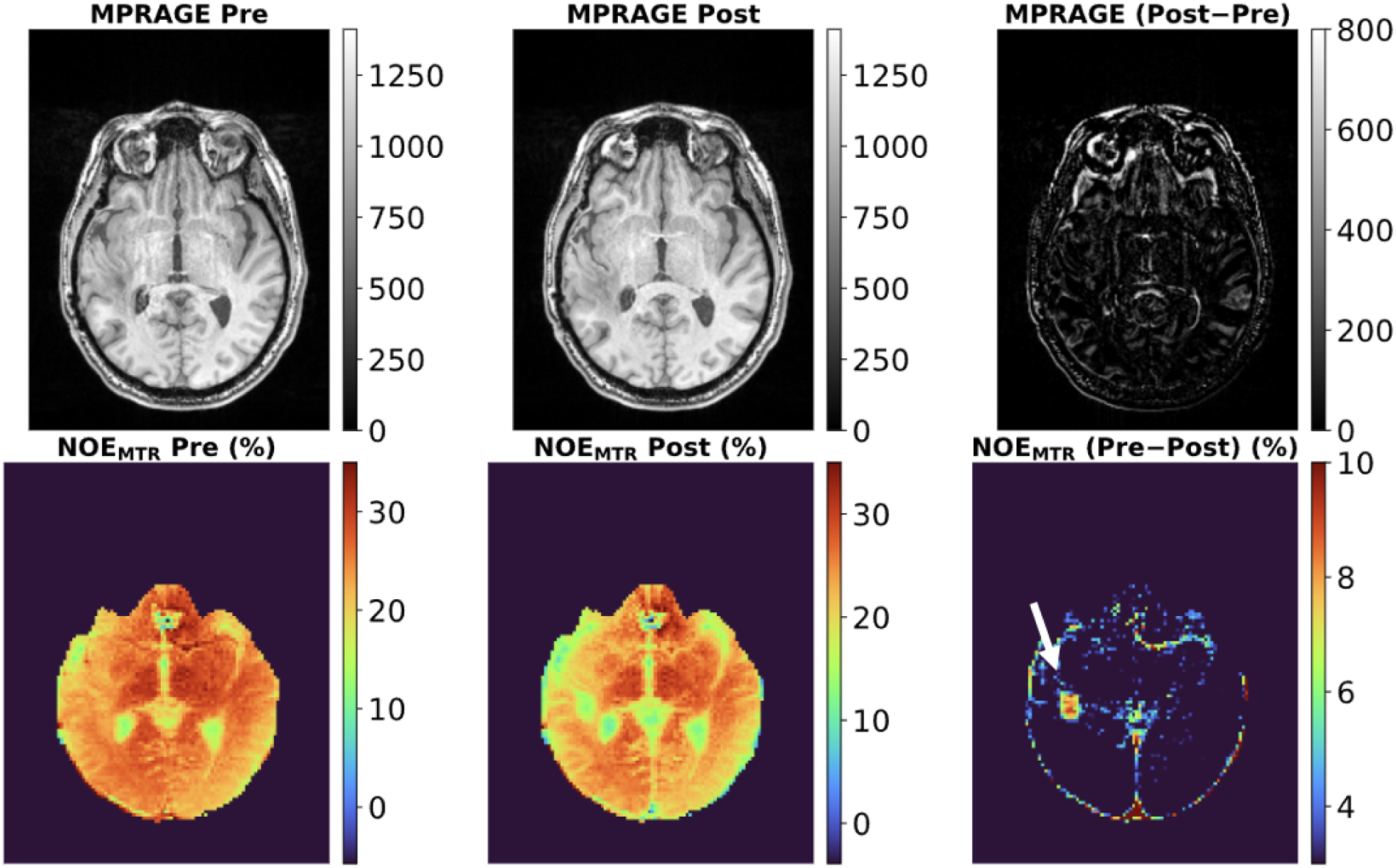
The first row contains MPRAGE (T1-weighted) images, which include pre- and post-Gd contrast, and a difference map from a human participant diagnosed with recurrent glioblastoma at 3T. The second row presents NOE_MTR_ maps pre- and post-Gd contrast, along with their difference map. Notably, no contrast enhancement is visible in the tumor regions on the post-Gd contrast MPRAGE images. However, the difference in the NOE_MTR_ map clearly highlights the tumor regions.

## Acknowledgements

Research reported in this publication was supported by the National Institute of Biomedical Imaging and Bioengineering of the National Institutes of Health under award number P41EB029460 and from the National Institute on Aging of the National Institutes of Health under award numbers RF1AG087306, R01AG063869, and R01AG091760.

